# Presynaptic depression maintains stable synaptic strength in developmentally arrested *Drosophila* larvae

**DOI:** 10.1101/840470

**Authors:** Sarah Perry, Pragya Goel, Daniel Miller, Barry Ganetzky, Dion Dickman

## Abstract

Positive and negative modes of regulation typically constrain synaptic growth and function within narrow physiological ranges. However, it is unclear how synaptic strength is maintained when both pre- and post-synaptic compartments continue to grow beyond stages imposed by typical developmental programs. To address whether and how synapses can adjust to a novel life stage for which they were never molded by evolution, we have characterized synaptic growth, structure and function at the *Drosophila* neuromuscular junction (NMJ) under conditions where larvae are terminally arrested at the third instar stage. While wild type larvae transition to pupae after 5 days, arrested third instar (ATI) larvae persist for up to 35 days, during which NMJs exhibit extensive overgrowth in muscle size, presynaptic release sites, and postsynaptic glutamate receptors. Remarkably, despite this exuberant growth of both pre- and post-synaptic structures, stable neurotransmission is maintained throughout the ATI lifespan through a potent homeostatic reduction in presynaptic neurotransmitter release. Arrest of the larval stage in *stathmin* mutants reveals a degree of progressive instability and neurodegeneration that was not apparent during the typical larval period. Hence, during a period of unconstrained synaptic growth through an extended developmental period, a robust and adaptive form of presynaptic homeostatic depression can stabilize neurotransmission. More generally, the ATI manipulation provides an attractive system for studying neurodegeneration and plasticity across longer time scales.

**SIGNIFICANCE STATEMENT:** It is unclear whether and how synapses adjust to a novel life stage for which they were never molded by evolution. We have characterized synaptic plasticity at the *Drosophila* neuromuscular junction in third instar larvae arrested in development for over 35 days. This approach has revealed that homeostatic depression stabilizes synaptic strength throughout the life of arrested third instars to compensate for excessive pre- and post-synaptic growth. This system also now opens the way for the study of synapses and degeneration over long time scales in this powerful model synapse.

## INTRODUCTION

Synapses are confronted with extensive challenges during development, maturation, and aging yet maintain stable information exchange. The dynamic and massive changes in synapse growth, pruning and remodeling, coupled with intrinsic adjustments in neuronal excitability can lead to unstable physiological activity. The resulting imbalances in excitation and inhibition would propagate within neural circuits to undermine network stability. To adapt to such challenges, synapses are endowed with the capacity to homeostatically adjust neurotransmission while still permitting the flexibility necessary for Hebbian forms of plasticity (Pozo and Goda, 2010; Prudencio et al., 2015; Turrigiano, 2012, 2017). The homeostatic control of neural activity operates throughout organismal lifespan to balance the tension between stability and flexibility, and is thought to break down in neurological and psychiatric diseases (Eichler and Meier, 2008; Hunt et al., 2017; Nelson and Valakh, 2015; Styr and Slutsky, 2018; Wondolowski and Dickman, 2013). Although it is clear that synapses have the capacity to express both Hebbian and homeostatic forms of plasticity, how these processes are integrated and balanced, particularly during development and aging, remain enigmatic.

The *Drosophila* larval neuromuscular junction (NMJ) is an accessible and versatile model for studying synaptic function, plasticity, and disease. This model glutamatergic synapse has enabled fundamental insights into synaptic growth, transmission, homeostatic plasticity, injury (Frank, 2014; Keshishian et al.; Li et al., 2018b; Menon et al., 2013). However, studies in this system are limited by the relatively short larval period of 3-4 days before pupariation, when NMJ accessibility is lost. This short temporal window limits the use of the third instar larval NMJ as a model for interrogating dynamic processes over chronic time scales. However, recent studies on the signaling cascades in *Drosophila* that control the transition from third instar to the pupal stage have revealed attractive targets for extending the duration of the third instar (Gibbens et al., 2011; Rewitz et al., 2009; Walkiewicz and Stern, 2009).

Developmental progression in *Drosophila* larvae is coordinated through two semi-redundant signaling pathways via Torso and insulin-like receptors that ultimately lead to ecdysone synthesis and release from the prothoracic gland (PG) to drive the transition from the larval stage to pupation (Rewitz et al., 2009; Walkiewicz and Stern, 2009; Yamanaka et al., 2013). A previous study reduced signaling through one arm of this pathway to extend the third instar stage from 5 to 9 days, where the important observation that NMJs continue to grow and function throughout this period was made (Miller et al., 2012). More recent work has demonstrated that loss of key transcription factors in the PG, including Smox (dSMAD2), can disrupt both signaling pathways to fully arrest larval development and prevent the transition to pupal stages (Gibbens et al., 2011; Ohhara et al., 2017) Remarkably, these arrested third instars (ATI) remain in the larval stage until death. The development of ATI larvae now provides an opportunity to characterize synaptic growth, function, and plasticity in a system of terminally persistent expansion beyond normal physiological ranges and has the potential to reveal new insights into processes such as neurodegeneration.

Here, we have developed an optimized approach to arrest *Drosophila* larvae at third instar stages to characterize NMJ growth, function, and plasticity. We find that ATI larvae continue to grow and survive for up to 35 days, where NMJs exhibit exuberant expansion in both pre- and post-synaptic compartments. Interestingly, this growth should enhance synaptic strength, yet no significant change is observed compared to baseline values. Instead, a potent reduction in presynaptic neurotransmitter release maintains stable synaptic strength across the life of an ATI larva. Finally, the ATI larvae enabled new insights into the progression of neurodegeneration in *stathmin* mutants. Together, arresting larval development now provides a powerful foundation to probe the mechanisms of synaptic growth, function, homeostatic plasticity, and neurodegeneration at a model glutamatergic synapse in a genetically tractable system.

## RESULTS

### Synaptic strength is maintained throughout the lifespan of an arrested third instar larva

To arrest larval development at the third instar stage, we targeted genes that could either disrupt both Torso and insulin signaling pathways or broadly inhibit the synthesis of ecdysone synthesis in the PG, processes ultimately necessary for the transition to pupal stages (Figure 1A; (Gibbens et al., 2011; Ohhara et al., 2017)). We reasoned that if we could prevent the release of ecdysone from the PG by knocking down a key transcript(s), pupation would be delayed indefinitely (Yamanaka et al., 2013). We screened several lines described by other investigators and found that a particular RNAi line targeting *smox* (dSMAD2), a transcription factor required for expression of both *torso* and insulin receptor genes, was the most effective, reliably preventing pupation in nearly all animals (Figure 1A-B). These developmentally arrested third instars (ATI) persist as larvae and live up to 35 days after egg lay (AEL). Typical wild type larvae spend ∼3 days in the third instar stage before pupation and metamorphosis, living beyond 60 days AEL as adults (Linford et al., 2013). For the first 5 days of development, ATI larvae appear largely unchanged compared to wild type, but they fail to progress to become “wandering” third instars. Rather, they continue to feed and gain body mass, peaking around 17 days AEL (ATI.17) and then gradually loosing body mass until dying soon after 33 days AEL (ATI.33) (Figure 1B). For further experiments, we compared wild type larvae at 5 days AEL (WT.5) to ATI larvae at varying time points, including 5 days AEL (ATI.5), a time point similar to wild type; 17 days AEL (ATI.17), a time corresponding to peak body mass; and 33 days AEL (ATI.33), a time near the terminal stage of the ATI lifespan.

**Figure 1:**
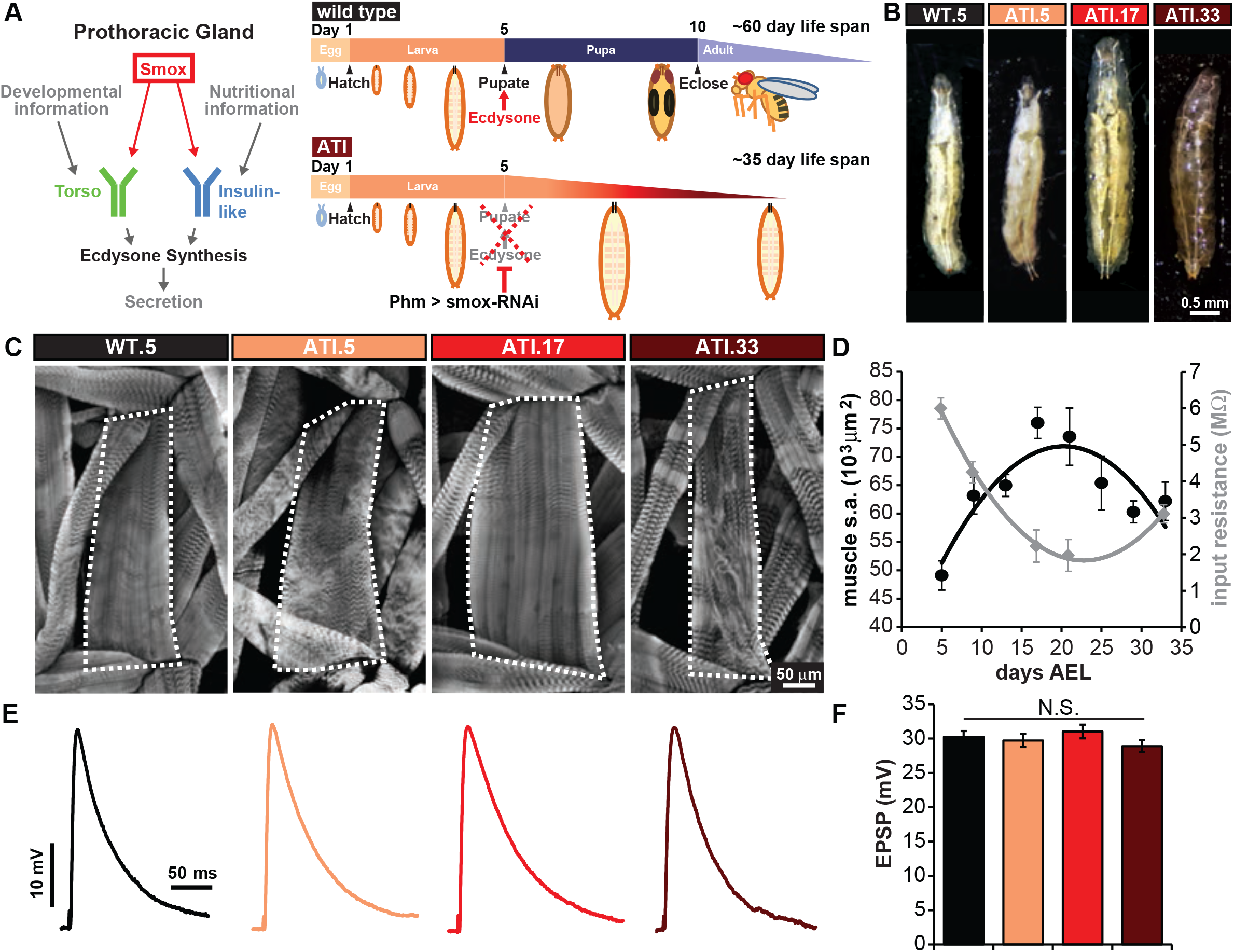
Synaptic strength remains stable throughout the life of an arrested third instar larvae. **(A)** (Left) Schematic illustrating the signaling pathway that stimulates ecdysone synthesis in the prothorasic gland prior to pupal formation. The transcription factor Smox is required for the expression of both Torso and insulin receptors. (Right) Schematic illustration comparing wild type fly development and *smox-RNAi* arrested development. **(B)** Representative photographs of third instar wild type larva (WT.5) and *smox-RNAi* (ATI) larva at different time points (days after egg lay (AEL)). **(C)** Representative images of larval body walls stained with anti-phalloidin to highlight muscle structure. M4 surface area is outlined in each image. **(D)** Graph summarizing muscle surface area measurements (black) and muscle compartment input resistance (grey) across ATI lifespan. **(E)** Representative EPSP traces for WT.5, ATI.5, 17 and 33 NMJs. **(F)** Average EPSP amplitudes for the genotypes shown in (E). Error bars indicate ±SEM. Additional details and statistical information (mean values, SEM, n, p, statistical tests data shown in F) are shown in Table 1-1.

To investigate NMJs across the ATI lifespan, we first characterized muscle size and passive electrical properties of the muscle. We observed a progressive gain in muscle size across the ATI lifespan, where muscle surface area increased by over 50%, peaking at ATI.17 and then decreasing to ATI.33 (Figure 1C-D). Consistent with this substantial increase in muscle size, electrophysiological recordings of NMJs across the ATI lifespan revealed a massive decrease in input resistance peaking around ATI.17 (Figure 1D). Remarkably, despite these changes in muscle size, synaptic strength (EPSP amplitude) remains constant across ATI NMJs (Figure 1E-F). Thus, as larvae grow and decline through an arrested third instar lifespan, synaptic strength at the NMJ remains constant.

### Presynaptic compartments at the NMJ progressively expand in ATI larvae

Clearly, ATI NMJs maintain synaptic strength despite the substantial increase in muscle size that progresses through arrested larval development. In principle, modulations to the number of presynaptic release sites (N), the probability of release at each individual release site (P_r_), and/or the postsynaptic response to glutamate release from single synaptic vesicles (quantal size, (Q)) could stabilize synaptic strength at these NMJs (Dittman and Ryan, 2019). We first assessed synaptic growth to determine whether the number of presynaptic release sites increases in proportion to the muscle surface area. During the conventional 3-4 day period of larval development, there is a 100-fold expansion in the NMJ, with changes to the passive electrical properties of the muscle and a concomitant growth of pre- and post-synaptic compartments (Atwood et al., 1993; Menon et al., 2013; Schuster et al., 1996). These changes are thought to scale NMJ function in parallel with growth and maintain sufficient depolarization for muscle contraction (Davis and Goodman, 1998). However, the progressive increase in muscle size at ATI NMJs poses a further challenge, where synapses may need to expand to compensate for overgrowth. We therefore considered whether adaptive changes in the growth of motor terminals and/or number of synapses served to stabilize synaptic strength (EPSP amplitude). Using immunostaining, we instead found a progressive enhancement in the neuronal membrane surface area and in the number of boutons per NMJ throughout the ATI lifespan (Figure 2A-D). In fact, the bouton to muscle area ratio even overshoots the scaling that is normally observed at conventional development between first and third instar larval stages (WT.5: 40 boutons/40,000 µm^2^ ratio (Schuster et al., 1996); ATI.17: 100 boutons/75,000 µm^2^ ratio (Table 1-1)). Hence, motor neuron terminals grow in excess to muscle growth.

**Figure 2:**
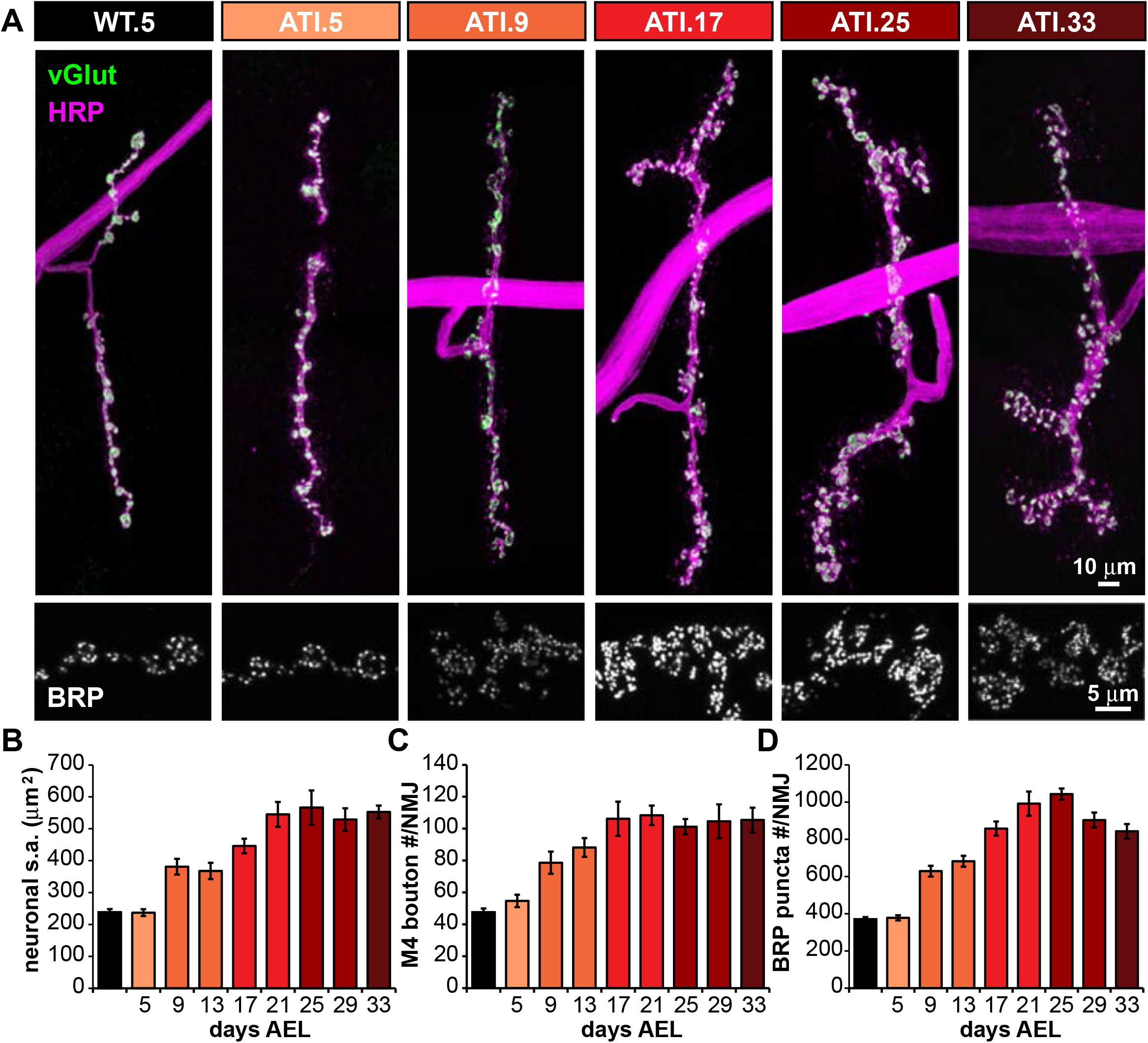
Progressive synaptic growth and a concomitant accumulation of release sites at ATI NMJs. **(A)** (Top) Representative M4-Ib images of NMJs for WT.5 and several ATI time points stained with anti-vGlut (vesicle-filled boutons, green) and anti-HRP (neuronal membrane, magenta). (Bottom) representative portion of the synapses above marked with anti-BRP (active zone scaffold). **(B-D)** Graphs showing the average neuronal membrane surface area (B), bouton number (C), and BRP puncta number (D) per muscle 4 NMJ for WT.5 and the indicated ATI time points. Error bars indicate ±SEM. Statistical information (mean values, SEM, n, p and additional BRP data) are shown in Table 1-1.

Since NMJ boutons expand across ATI stages, we considered the possibility of a compensatory reduction in the number of release sites. There is precedence for a reduction in the density of active zones (AZs), independent of NMJ growth, to maintain synaptic strength (Graf et al., 2009). To identify individual presynaptic release sites, we immunostained NMJs with an antibody against Bruchpilot (BRP), a central scaffolding protein that constitutes the “T-bar” structure at AZs in *Drosophila* (Kittel et al., 2006; Wagh et al., 2006). Since ∼96% of release sites are labeled by BRP at the fly NMJ (Akbergenova et al., 2018; Gratz et al., 2019; Wagh et al., 2006), we defined a release site as a BRP punctum and quantified these structures across the ATI lifespan. Interestingly, we found no significant changes in BRP puncta density across ATI stages, with total BRP puncta number per NMJ increasing in proportion to neuronal membrane area and bouton number (Figure 2A-D, Table 1-1). Finally, although the number of BRP puncta increased, the size and fluorescence intensity of these puncta can be reduced at NMJs to compensate for synaptic overgrowth, reducing Pr at individual release sites and maintaining overall synaptic strength (Goel et al., 2019a, 2019b). However, although BRP number at ATI NMJs increases to over three-fold that of wild-type NMJs, no compensatory reduction in size and/ or intensity of BRP puncta was observed (Figure 2A lower panel, Table 1-1). Indeed, BRP puncta intensity was significantly *increased* compared to WT.5 levels (Table 1-1), which may reflect the age-dependent increase size and intensity and active zones documented at the fly NMJ (Akbergenova et al., 2018). Thus, this anatomical analysis reveals an increase in the number of release sites (N) at ATI NMJs, implying that other adaptations compensate for excessive growth in ATI larvae.

### Postsynaptic receptor fields accumulate at the NMJ over the ATI lifespan

Given the substantial increase in AZ number and intensity but stable synaptic strength, we next considered the possibility that a reduction in the postsynaptic receptivity to neurotransmitter (Q) may have served to offset the observed presynaptic overgrowth at ATI NMJs. One possibility is that a reduction in the abundance, composition, and/or function of postsynaptic glutamate receptors (GluRs) may have occurred at ATI NMJs. At the fly NMJ, two receptor subtypes containing either GluRIIA- or GluRIIB-subunits form complexes with the essential GluRIIC, GluRIID, and GluRIIE subunits to mediate the postsynaptic currents driving neurotransmission (DiAntonio, 2006; Qin et al., 2005). GluRIIA-containing receptors mediate larger current amplitudes and slower decay kinetics compared to the GluRIIB-containing receptor counterparts (Han et al., 2015; Petersen et al., 1997). We examined the postsynaptic GluRs using antibodies that specifically recognize the GluRIIA- or GluRIIB-subunits, as well as the common GluRIID subunit. Consistent with presynaptic overgrowth, total GluR puncta numbers per NMJ mirrored the increase in presynaptic AZ number (Figure 3B). Similarly, we observed a significant increase in the abundance of all GluR subunits assessed at ATI NMJs revealed by enhanced fluorescence intensity (Figure 3C). Together, this demonstrates that postsynaptic receptor fields progressively expand in number and abundance, mirroring the accumulation in presynaptic structures across the ATI lifespan.

**Figure 3:**
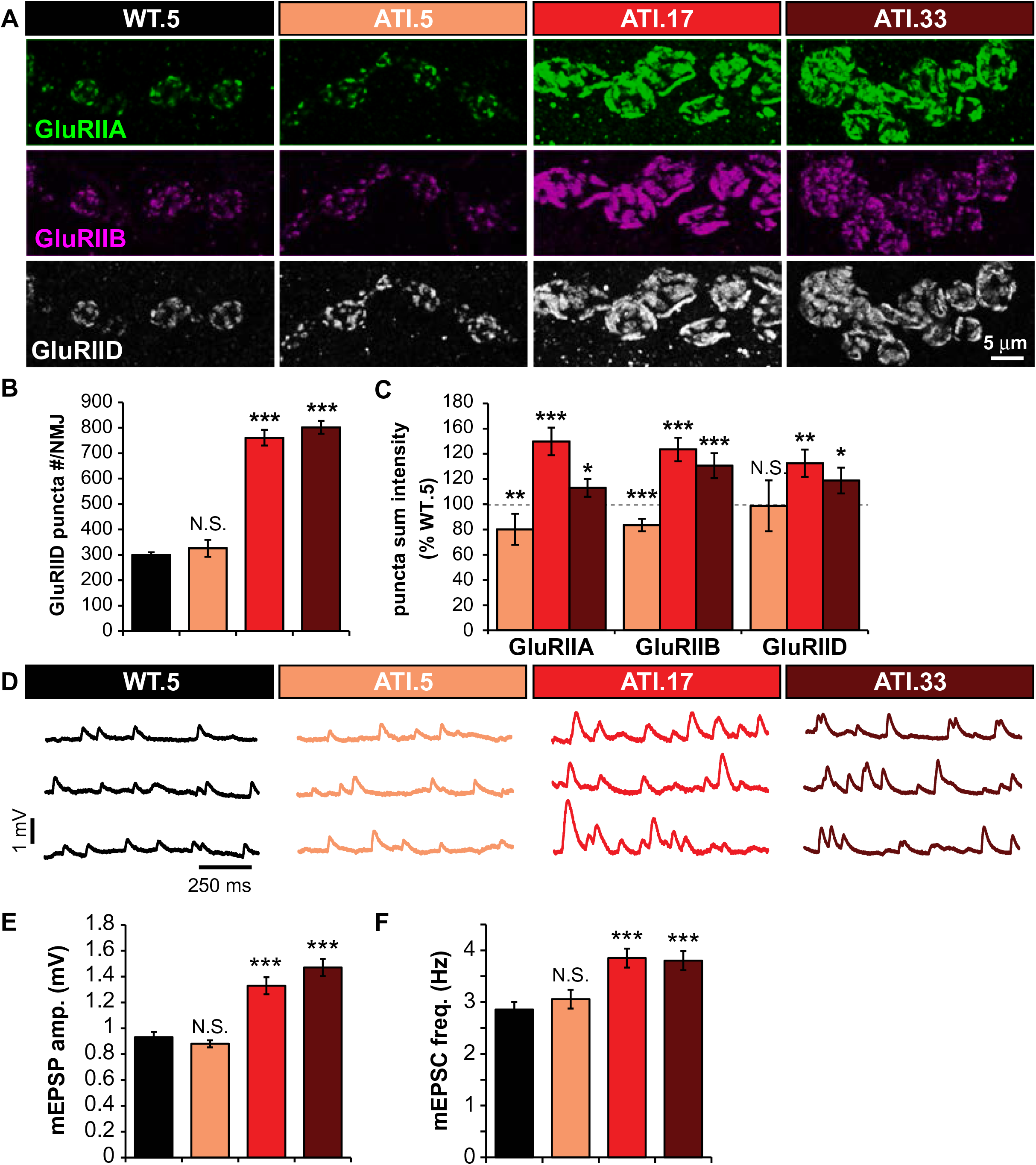
Postsynaptic glutamate receptors accumulate and quantal size increases over the ATI lifespan. **(A)** Representative images of the indicated GluR subunit staining at NMJ terminals of muscle 4 (1b boutons) in wild type (WT.5) and the indicated ATI time points. Quantification of GluRIID puncta number **(B)** and GluR puncta sum intensity **(C)** in the indicated genotypes. **(D)** Representative mEPSP traces of WT.5 and the indicated ATI time points. Quantification of mEPSP amplitude **(E)** and frequency **(F)** in genotypes shown in (D). Error bars indicate ±SEM. *p≤0.05; **p≤0.01; ***p≤0.001; N.S = not significant, p>0.05. Detailed statistical information (mean values, SEM, n, p) is shown in Table 1-1.

We next considered whether an apparent reduction in GluR functionality compensated for the expansion of glutamate receptor fields at ATI NMJs. We determined GluR functionality by electrophysiologically recording miniature events at ATI NMJs. Consistent with the increased fluorescence intensity of all subunits, we observed an ∼50% increase in mEPSP amplitude compared to wild-type levels in ATI.17 larvae, an enhancement that persisted through ATI.33 (Figure 3D-E). Consistent with increased presynaptic growth, we also observed an increase in mEPSP frequency (Figure 3F). Increased expression of GluRIIA-containing receptors at the fly NMJ enhances mEPSP amplitude, as expected, but does not alter presynaptic neurotransmitter release (Li et al., 2018c; Petersen et al., 1997). Thus, the increase in both AZ number (N) and quantal size (Q) at ATI NMJs should have together elicited a larger evoked response. However, EPSP amplitude is unchanged throughout the ATI lifespan, implying that a reduction in release probability (P_r_) of sufficient magnitude must be induced to fully counteract the increase in N and Q to maintain stable synaptic strength at ATI NMJs.

### Synaptic strength at ATI NMJs is maintained through a potent homeostatic decrease in release probability

ATI larvae exhibit exuberant synaptic growth with accumulations of both pre- and post-synaptic components, resulting in an increased N and Q, factors that should enhance synaptic strength. However, EPSP amplitudes remain stable across the ATI lifespan, implying presynaptic release probability (P_r_) must be substantially and precisely diminished to compensate. To further test this idea, we calculated quantal content (the number of synaptic vesicles released per stimulus) and found a substantial reduction at ATI NMJs (Figure 4A). Next, we assessed presynaptic function independently of mEPSP amplitude by performing failure analysis, where repeated stimulations in low extracellular Ca^2+^ (0.15 mM) fail to elicit a response in ∼50% of trials in wild type. At ATI NMJs, failure rate was markedly increased (Figure 4B), consistent with reduced quantal content. Finally, we assayed paired-pulse ratios to gauge P_r_. At low extracellular Ca^2+^ (0.3 mM), paired-pulse facilitation (PPF) is observed at wild-type NMJs, while paired-pulse depression (PPD) is found in elevated Ca^2+^ (1.5 mM)(Böhme et al., 2016; Li et al., 2018c). In ATI.17 and ATI.33 NMJs, PPF was significantly increased while PPD was reduced, consistent with reduced P_r_ relative to wild type (Figure 5C-D). It is interesting to note that a similar phenomenon has been observed at the *Drosophila* NMJ in the context of typical larval development, referred to as presynaptic homeostatic depression (PHD). Here, mEPSP size is enhanced while quantal content is reduced to maintain normal EPSP amplitudes (Daniels et al., 2004; Gaviño et al., 2015; Li et al., 2018c). While it is not clear that the mechanism of depression is shared between later ATI time points and PHD, we can posit that a homeostatic reduction in presynaptic release probability compensates for increased quantal size to maintain synaptic strength across the ATI life span.

**Figure 4:**
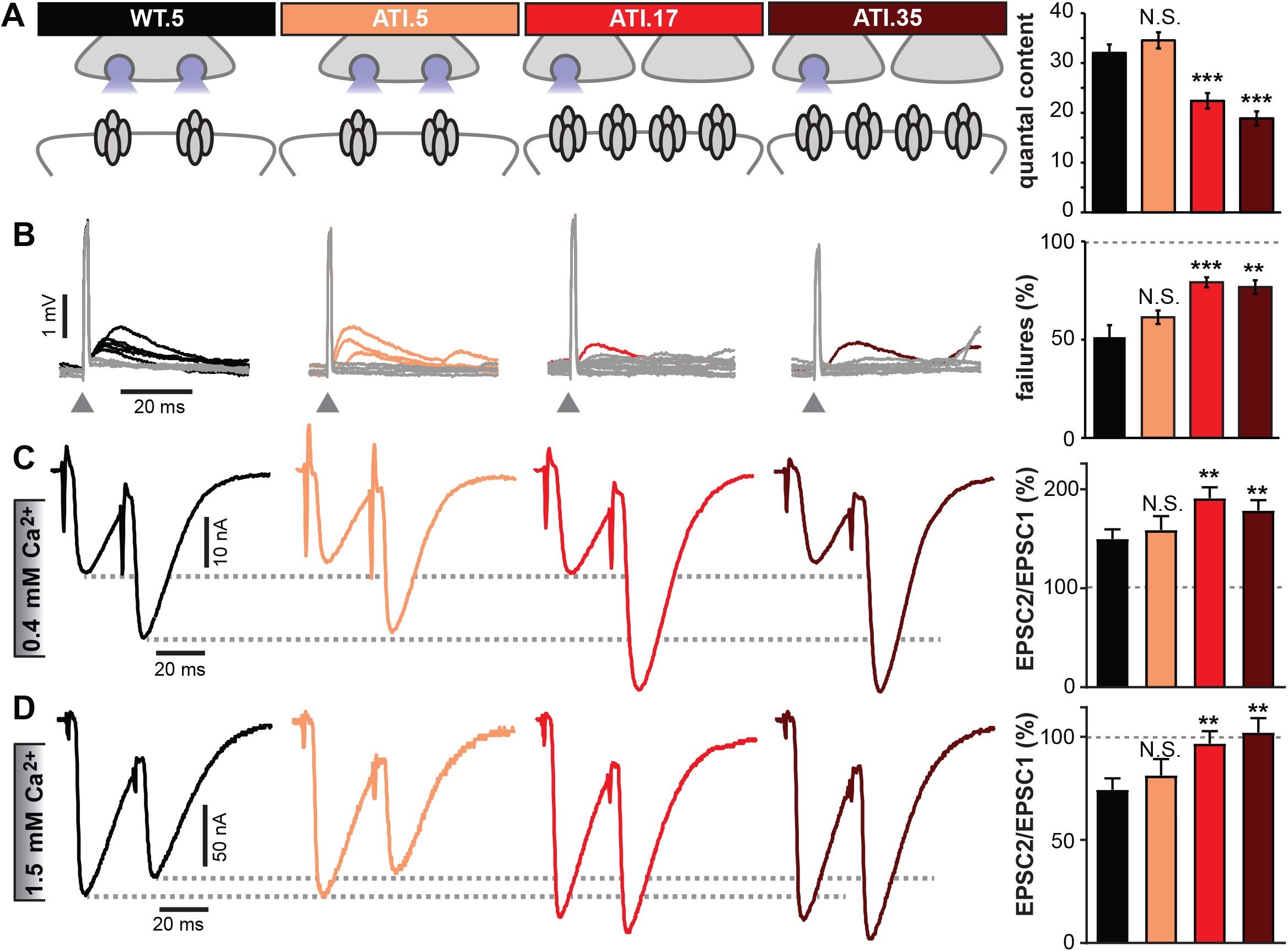
A potent reduction in neurotransmitter release probability is expressed across the ATI lifespan. **(A)** (Left) Schematic in the indicated genotypes illustrating reduced synaptic strength at later ATI time points. (Right) Quantal content calculated from EPSP and mEPSP data in Figs. 1 and 3. **(B)** (Left) Representative traces of attempted stimulations during failure analysis. Grey traces indicate failures and colored traces indicate successful evoked responses. Eight traces are shown for each genotype. (Right) Quantification of % failures for each genotype. **(C)** Representative two electrode voltage clamp (TEVC) traces for showing paired pulse facilitation for each genotype (left) and quantification of paired pulse ratio (right). **(D)** Representative TEVC traces for paired pulse depression for each genotype (left) and quantification of paired pulse ratio (right). Error bars indicate ±SEM. **p≤0.01; ***p≤0.001; N.S. = not significant, p>0.05. Detailed statistical information (mean values, SEM, n, p) is shown in Table 1-1.

**Figure 5:**
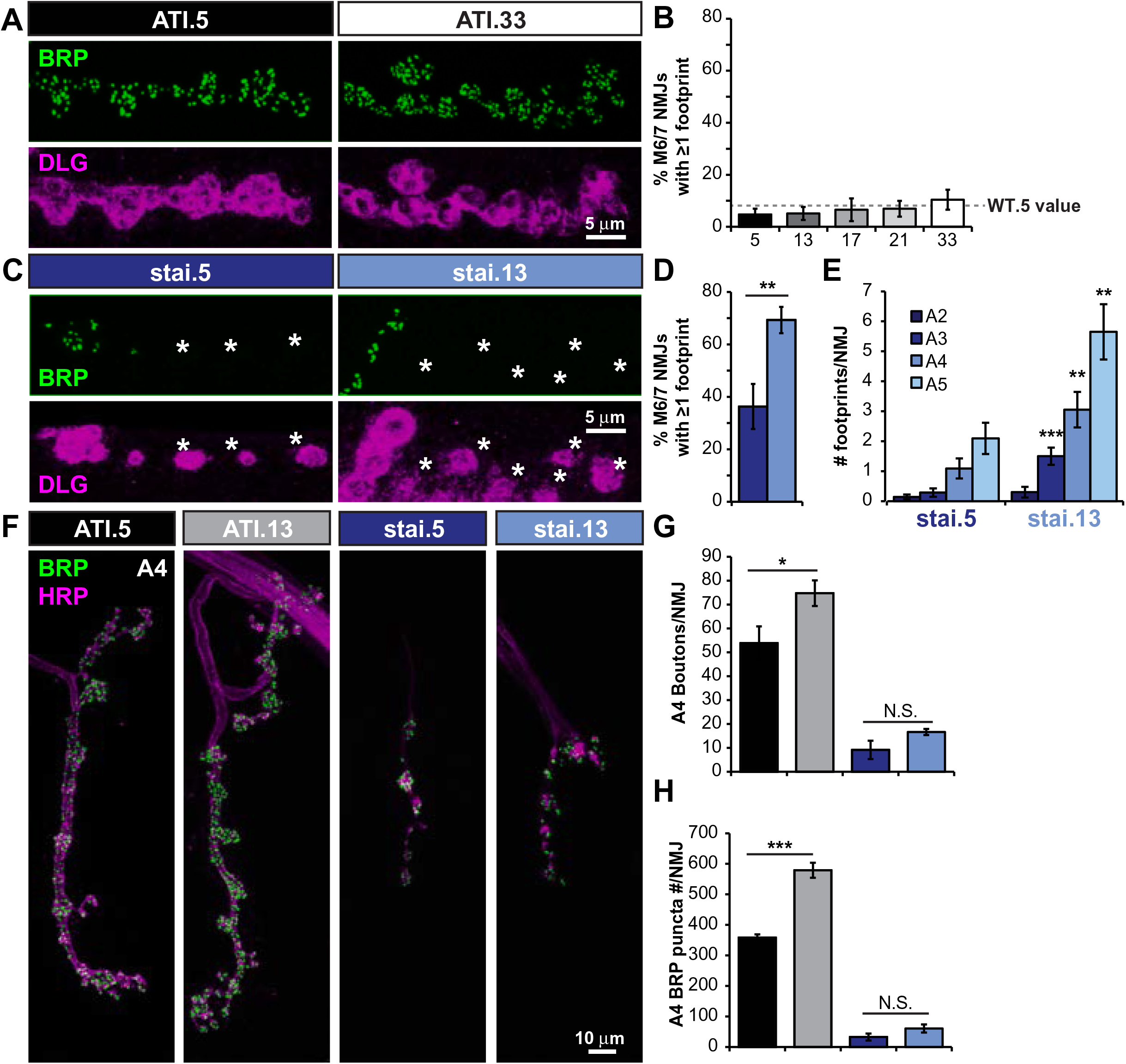
Extending the larval stage reveals the progression of synaptic retractions in *stathmin* mutants. **(A)** Representative images of ATI.5 and ATI.33 synapses stained with presynaptic (BRP) and postsynaptic (DLG) markers demonstrating a lack of presynaptic retractions at these stages. **(B)** Quantification of % NMJs at muscle 6/7 with one or more footprint observed across the ATI lifespan. The value for wild type at day 5 is shown (WT.5) by a dashed line. **(C)** Representative BRP and DLG images of *stathmin* mutant NMJs in an ATI background (stai.5 and stai.13, see Table 1-1 for full genotypes) showing footprints (DLG staining without corresponding BRP marked with an asterisk). **(D)** Quantification of % NMJs with one or more footprints in stai.5 and stai.15 animals. **(E)** Quantification of footprints per NMJ by segment in *stathmin* mutants demonstrating more severe effects on posterior segments over time. **(F)** Representative images of wild type and *stathmin* ATI NMJs at muscle 4 (Ib boutons) at day 5 and 13 stained with HRP (neuronal membrane) and BRP (active zone scaffold) showing a failure of synaptic growth in *stai* mutants. Quantification of bouton number **(G)** and BRP puncta number **(H)** per NMJ on segment A4 for the indicated genotypes. Error bars indicate ±SEM. *p≤0.05; **p≤0.01; ***p≤0.001; N.S.=not significant, p>0.05. Detailed statistical information (mean values, SEM, n, p) is shown in Table 1-1.

It has previously been shown that NMJs expressing PHD can also express other forms of homeostatic plasticity, including a process referred to as presynaptic homeostatic potentiation (PHP) (Gaviño et al., 2015; Goel et al., 2019a; Li et al., 2018c). To induce PHP, we applied the postsynaptic GluR antagonist philanthotoxin-343 (PhTx) (Frank et al., 2006). 10 min incubation in PhTx reduces mEPSP amplitude in both wild type and ATI NMJs, as expected (Figure 4-1). In turn, EPSP amplitude is maintained at baseline levels due to a retrograde, homeostatic increase in presynaptic neurotransmitter release in wild type. Similarly, PHP is robustly expressed across ATI NMJs (Figure 4-1). Thus, like PHD and other forms of homeostatic plasticity studied at the *Drosophila* NMJ, the presynaptic inhibition observed at ATI NMJs can be balanced with acute GluR challenge to express PHP and maintain stable synaptic strength.

### Extending the larval stage reveals the progression of axonal degeneration in *stathmin* mutants

In our final set of experiments, we considered whether ATI larvae could be utilized as models for aging and/or neurodegeneration. We hypothesized that NMJs in ATI larvae were unlikely to exhibit classical hallmarks of aging synapses. Although muscle integrity appears to degrade in ATI.33 compared to earlier time points (Figure 1), synaptic growth (Figure 2), GluR receptor fields (Figure 3), and presynaptic function (Figure 4) all appear similar in ATI.33 relative to earlier time points. Indeed, while reductions in synaptic components and neurotransmission have been observed at aging mammalian NMJs (Li et al., 2018a; Taetzsch and Valdez, 2018), NMJ structure and function remains surprisingly robust in ATI larvae nearing death, with no apparent defects in synaptic function or even PHP plasticity. One additional canonical indicator of aging reported at mammalian NMJs includes presynaptic retractions and fragmentation (Li et al., 2018a; Taetzsch and Valdez, 2018). We therefore assessed synaptic retractions across the ATI lifespan using an established “footprint” assay, in which a postsynaptic marker is observed to persist without a corresponding presynaptic marker (Eaton et al., 2005; Graf et al., 2011; Perry et al., 2017). However, ATI NMJs, including ATI.33, showed surprisingly stable synapses, with no apparent increases in footprints compared with earlier time points (Figure 5A-B). Together, these results indicate that NMJ structure, function, and integrity remain surprisingly robust across all stages of ATI larvae, even at terminal periods, and are therefore unlikely to serve as a compelling model for age-related synaptic decline.

Although NMJs remain structurally intact and stable across the lifespan of the ATI larvae, this manipulation does enable a substantially longer time scale compared to the typical 5 days of larval development to investigate insults contribute to neuronal degeneration. We chose to characterize NMJ growth and stability in *stathmin* mutants extended through the ATI manipulation. Stathmin is a tubulin-associated factor involved in maintaining the integrity of the axonal cytoskeleton (Duncan et al., 2013; Graf et al., 2011; Lachkar et al., 2010; Ozon et al., 2002). The mammalian homolog of *Drosophila stathmin* (*SCG10*) is highly conserved and is thought to function as a surveillance factor for axon damage and degenerative signaling (Shin et al., 2014). In *Drosophila*, loss of *stathmin* leads to a marked increase in NMJ footprints, where more posterior segments show increased severity relative to more anterior segments (Graf et al., 2011). Surprisingly, *stathmin* mutants are still able to pupate and develop into adults. However, *stathmin* mutants extended in larval stages by the ATI manipulation die shortly after 21 days AEL. We therefore sought to use the ATI system to determine the impact of a prolonged phase of axonal instability in *stathmin* mutants. Indeed, NMJs exhibit increased footprints in stai.13 (*stathmin* mutants extended to ATI.13 time points) when compared to stai.5 controls in both frequency (Figure 5C-D) and severity (Figure 5E), with the most severe retractions observed in posterior abdominal segments (A3-A5). Finally, we tested whether NMJ growth increased in ATI-extended *stathmin* NMJs, as it does in wild type. While control ATI synapses grow in bouton and BRP puncta number between 5 and 13 days AEL, *stathmin* NMJs fail to consistently expand (Figure 5F-H). These experiments highlight the potential of the ATI system to be a useful tool for defining the progression of neurodegeneration at the *Drosophila* NMJ, which is otherwise limited to short larval stages.

## DISCUSSION

By arresting further maturation at third instar *Drosophila* larvae, we have been able to accomplish a detailed study of NMJ structure, function, and plasticity over much longer timescales than previously possible. This ATI larval system has revealed how the NMJ maintains stable transmission over a vastly extended developmental timescale, where persistent overgrowth in both pre- and post-synaptic compartments is offset through a potent and homeostatic reduction in neurotransmitter release. Hence, this study not only provides evidence for a potentially novel homeostatic signaling system that balances release probability with synaptic overgrowth but now extends the temporal window to enable the characterization of a variety of processes, including neurodegeneration, at a powerful model synapse.

As described by Miller et al. (2012), NMJs in third instar larvae that have been developmentally arrested for at least a week beyond the normal time of pupariation continue to grow and add new boutons. Here we extend this observation to larvae arrested at the third instar for over 30 days, further demonstrating that mechanisms do not exist to suppress or negatively regulate growth when developmental timing is artificially extended. During normal larval growth from first to third instar, the body wall muscles undergo rapid and immense expansion, growing nearly 100-fold in surface area within a few days (Atwood et al., 1993; Menon et al., 2013). Presynaptic terminals grow and add new boutons in parallel with muscle growth, presumably to maintain stable NMJ strength. In effect, sufficient levels of muscle excitation is sustained through a coordinated increase in all three parameters controlling synaptic physiology: N (number of release sites), P (release probability at each site), and Q (quantal size) (Neher, 2015) Hence, during typical stages of larval development, increasing muscle growth requires a concomitant elaboration in NMJs, implying robust signaling systems exist to ensure synaptic size, structure, and function expand in a coordinated manner. This tight structural coupling between muscle fiber and NMJ growth is also observed in mammals and is thought to be a primary mechanism for maintaining NMJ strength during post-developmental muscle growth or wasting (Balice-Gordon et al., 1990; Sanes and Lichtman, 1999, 2001). However, when the normal developmental program is made to continue without terminating in pupariation, NMJ growth continues apparently unchecked, posing a potential challenge of hyperexcitation. There is emerging evidence that when NMJ growth is genetically perturbed, a redistribution of active zone material or adaptations in synapse morphogenesis or postsynaptic neurotransmitter receptors can maintain stable synaptic strength (Bae et al., 2016; Goel et al., 2019a, 2019b; Graf et al., 2009). In the case of NMJ overgrowth in *endophilin* mutants, a homeostatic scaling in active zone size compensates for increased number to lower release probability and maintain stable synaptic strength (Goel et al., 2019a). However, NMJs in ATI larvae do not appear to utilize such strategies. Rather, a latent form of adaptive plasticity is revealed at ATI NMJs that is sufficiently potent and precise to inhibit neurotransmitter release probability and compensate for the overgrowth of both pre- and post-synaptic compartments.

The presynaptic inhibition of neurotransmitter release that maintains synaptic strength at ATI NMJs is a potentially novel phenomenon of homeostatic plasticity. This form of presynaptic depression appears to be an entirely functional change that reduces release probability, without any apparent adaptations to active zone number, intensity or synaptic structure. Electrophysiologically, the presynaptic inhibition demonstrated at ATI NMJs resembles presynaptic homeostatic depression (PHD), a form of homeostatic plasticity characterized at the *Drosophila* NMJ in which excess glutamate release induces a compensatory reduction in release probability that maintains stable synaptic strength (Daniels et al., 2004; Gaviño et al., 2015; Li et al., 2018c). Like PHD, the presynaptic inhibition at ATI NMJs is not reflected in changes to the active zones or synaptic structure (Goel et al., 2019a; Gratz et al., 2019; Li et al., 2018c). However, the only mechanisms known to be capable of inducing PHD require enhanced synaptic vesicle size that results from endocytosis mutants or overexpression of the vesicular glutamate transporter (Daniels et al., 2004; Dickman et al., 2005; Goel et al., 2019a; Verstreken et al., 2002; Winther et al., 2013). In contrast, there is no evidence for changes in synaptic vesicle size at ATI NMJs, as the enhanced postsynaptic glutamate receptor levels observed are sufficient to explain the increased quantal size (Fig. 3). Hence, if the homeostatic depression observed at ATI NMJs is ultimately the same plasticity mechanism as PHD, then this would be the first condition that does not require enlarged synaptic vesicle size. In this case, perhaps excess global glutamate release from increased release sites at ATI NMJs induces the same homeostatic plasticity that increased glutamate released from individual synaptic vesicles does. This would be consistent with a “glutamate homeostat”, responding to excess presynaptic glutamate release, necessary to induce and express PHD (Li et al., 2018c). Alternatively, the presynaptic inhibition triggered at ATI NMJs could be a novel form of presynaptic homeostatic depression which is induced in response to synaptic overgrowth. Interestingly, while increased postsynaptic glutamate receptors levels enhance mini size, no adaptive change in presynaptic function results, which leads to a concomitant increase in synaptic strength (DiAntonio et al., 1999; Li et al., 2018c). One possibility is that a coordinated increase in both pre- and post-synaptic compartments may be necessary to induce the presynaptic inhibition observed at ATI NMJs. The ATI model provides a unique opportunity to interrogate the interplay between developmental growth, adaptive presynaptic inhibition, and other homeostatic signaling systems.

Extending the larval stage through the ATI manipulation will circumvent limitations of the brief time window provided by the standard developmental program. Although the ATI model does not appear to exhibit the features described at aging mammalian NMJs (Li et al., 2018a; Taetzsch and Valdez, 2018), we have demonstrated its potential for modeling neurodegenerative conditions by showing the extent of synaptic destabilization caused by loss of *stathmin* that was not fully apparent when restricted to the normal short developmental period in *Drosophila* larvae (Graf et al., 2011). In particular, by examining *stathmin* mutant phenotypes in ATI-extended larvae, we were able to observe progressive, time-dependent retractions of presynaptic terminals and gain further insight into *stathmin’s* role in normal NMJ growth and stability. Consistent with the role of *stathmin* in flies, the mammalian homolog (*SCG10*) is thought to be part of an axonal injury surveillance system, where it accumulates after injury and is involved in regenerative signaling (Shin et al., 2014). More generally, previous studies of degenerative disease models in the larval system have been limited by the brief timespan. For example, one important ALS disease model in flies involves overexpression of repetitive RNAs and peptides derived from the human *C9ORF72* gene (Mizielinska et al., 2014; Xu et al., 2013). However, while a variety of progressive and degenerative phenotypes are observed in photoreceptors of adult flies, only the most toxic transgenes are capable of inducing substantial neurodegeneration at the larval NMJ (Perry et al., 2017), likely due to the limited time frame of typical larval development. The longer timescale enabled by the ATI model therefore provides new opportunities to study progressive phenotypes during neuronal injury, stress, and neurodegeneration in addition to the plasticity discussed above in a rapid and genetically tractable system. Indeed, fly models of neurodegenerative conditions such as ALS, Huntington’s, Parkinson’s and Alzheimer’s diseases (McGurk et al., 2015) can benefit from the high resolution imaging and electrophysiological approaches established at the larval NMJ. The powerful combination of established genetic tools, including binary expression systems (Gal4/UAS, LexA, QF systems; (Venken et al., 2011)) and emerging CRISPR/Cas9 manipulations (Bier et al., 2018) with the ATI model provides an exciting foundation to gain new insights into synaptic growth, structure, function, plasticity, injury, and neurodegeneration over long times using the glutamatergic NMJ as a model.

## MATERIALS AND METHODS

### Fly Stocks

*Drosophila* stocks were raised at 25°C on standard molasses food. The *w^1118^* strain is used as the wild type control unless otherwise noted, as this is the genetic background of the genetic mutants used in this study. ATI larvae were generated by crossing *phm-GAL4* (Miller et al., 2012) to *UAS*-*smox-RNAi* (BDSC #41670). *Stathmin* mutations were introduced into the ATI background (*stai* allele: BDSC #16165). All experiments were performed on male or female third-instar larvae or arrested third instar larvae at various time points. A complete list of all stocks and reagents used in this study, see Table 2-1.

### Immunocytochemistry

Third-instar male or female larvae were dissected in ice cold 0 Ca^2+^ HL-3 and fixed in Bouin’s fixative for 5 min as described (Chen et al., 2017). Briefly, larvae were washed with PBS containing 0.1% Triton X-100 (PBST) for 30 min, blocked for an hour with 5% normal donkey serum in PBST, and incubated overnight in primary antibodies at 4°C followed by washes and incubation in secondary antibodies. Samples were mounted in VectaShield (Vector Laboratories). The following antibodies were used: mouse anti-Bruchpilot (nc82; 1:100; Developmental Studies Hybridoma Bank; DSHB); rabbit anti-DLG ((1:10,000; (Pielage et al., 2005)); guinea pig anti-vGlut ((1:2000; (Goel and Dickman, 2018)); mouse anti-GluRIIA (8B4D2; 1:100; DSHB); affinity purified rabbit anti-GluRIIB (1:1000; (Perry et al., 2017)), guinea pig anti-GluRIID ((1:1000; (Perry et al., 2017)). Donkey anti-mouse, anti-guinea pig, and anti-rabbit Alexa Fluor 488-, Cyanine 3 (Cy3)-, and Dy Light 405-conjugated secondary antibodies (Jackson Immunoresearch) were used at 1:400. Alexa Fluor 647 conjugated goat anti-HRP (Jackson ImmunoResearch) was used at 1:200. All antibody information is summarized in Table 2-1.

### Confocal imaging and analysis

Samples were imaged using a Nikon A1R Resonant Scanning Confocal microscope equipped with NIS Elements software and a 100x APO 1.4NA oil immersion objective using separate channels with four laser lines (405, 488, 561, and 637 nm). For fluorescence quantifications of BRP intensity levels, z-stacks were obtained using identical settings for all genotypes with z-axis spacing 0.5 µm within an experiment and optimized for detection without saturation of the signal as described (Perry et al., 2017). Boutons were counted using vGlut- and HRP-stained Ib NMJ terminals on muscle 4 of segment A2-A4, considering each vGlut punctum to be a bouton. The general analysis toolkit in the NIS Elements software was used for image analysis as described (Kikuma et al., 2017). Neuronal surface area was calculated by creating a mask around the HRP channel that labels the neuronal membrane. BRP puncta number, area, and total BRP intensity per NMJ were quantified by applying by using a bright-spot detection method and filters to binary layers on the BRP labeled 488 channel in a manner similar to that previously described (Goel et al., 2019b). GluRIIA, GluRIIB, and GluRIID puncta intensities were quantified by measuring the total sum intensity of each individual GluR punctum and these values were then averaged per NMJ to get one sample measurement (n). For NMJ retraction analysis, footprints were scored by eye as reported in (Eaton et al., 2005) on M6/7 segments A2-A5. Anti-DLG was used as a postsynaptic marker and either anti-vGlut or anti-BRP for a presynaptic marker (wild type controls yielded similar retraction scores for either presynaptic marker).

### Electrophysiology

All dissections and recordings were performed in modified HL-3 saline (Stewart et al., 1994; Dickman et al., 2005; Kiragasi et al., 2017) containing (in mM): 70 NaCl, 5 KCl, 10 MgCl_2_, 10 NaHCO_3_, 115 Sucrose, 5 Trehelose, 5 HEPES, and 0.4 CaCl_2_, pH 7.2. Neuromuscular junction sharp electrode (electrode resistance between 10-30 MΩ) recordings were performed on muscles 6 and 7 of abdominal segments A2 and A3 in wandering third-instar larvae as described (Goel et al., 2019a). Recordings were performed on an Olympus BX61 WI microscope using a 40x/0.80 water-dipping objective, and acquired using an Axoclamp 900A amplifier, Digidata 1440A acquisition system and pClamp 10.5 software (Molecular Devices). Electrophysiological sweeps were digitized at 10 kHz and filtered at 1 kHz. Data were analyzed using Clampfit (Molecular devices), MiniAnalysis (Synaptosoft), and Excel (Microsoft) software.

Miniature excitatory postsynaptic potentials (mEPSPs) were recorded in the absence of any stimulation and cut motor axons were stimulated to elicit excitatory postsynaptic potentials (EPSPs). Average mEPSP, EPSP, and quantal content were calculated for each genotype by dividing EPSP amplitude by mEPSP amplitude. Muscle input resistance (R_in_) and resting membrane potential (V_rest_) were monitored during each experiment. Recordings were rejected if the V_rest_ was above −60 mV, if the R_in_ was less than 5 MΩ, or if either measurement deviated by more than 10% during the course of the experiment. Larvae were incubated with or without philanthotoxin-433 (PhTx; Sigma; 20 μM) resuspended in HL-3 for 10 mins, as described (Frank et al., 2006; Dickman and Davis, 2009).

Failure analysis was performed in HL-3 solution containing 0.15 mM CaCl_2_, which resulted in failures in about half of the stimulated responses in wild-type larvae. A total of 40 trials (stimulations) were performed at each NMJ in all genotypes. Failure rate was obtained by dividing the total number of failures by the total number of trials (40). Paired-pulse recordings were performed at a Ca^2+^ concentration of 0.3 mM to assay facilitation (PPF) and 1.5 mM for depression (PPD). Following the first AP stimulation, a second EPSC was evoked at an interstimulus interval of 16.67 ms (60 Hz). Paired-pulse ratios were calculated as the EPSC amplitude of the second response divided by the first response (EPSC2/EPSC1).

### Experimental Design and Statistical Analysis

For electrophysiological and immunostaining experiments, each NMJ terminal (muscle 6 for physiology, and muscle 4 for immunostaining analyses) is considered an n of 1 since each presynaptic motor neuron terminal is confined to its own muscle hemi-segment. For these experiments, muscles 4 or 6 were analyzed from hemi-segments A2-A4 from each larvae, typically 2 NMJs/animal per experiment. To control for variability between larvae within a genotype, NMJs were analyzed from at least 5 individual larvae. See Table 1-1 for additional details.

Statistical analysis was performed using GraphPad Prism (version 7.0) or Microsoft Excel software (version 16.22). Data were assessed for normality using a D’Agostino-Pearson omnibus normality test, which determined that the assumption of normality of the sample distribution was not violated. Normally distributed data were analyzed for statistical significance using a Student’s t-test (pairwise comparison) or an analysis of variance (ANOVA) and Tukey’s test for multiple comparisons. Data were then compared using either a one-way ANOVA and tested for significance using a Tukey’s multiple comparison test or using an unpaired 2-tailed Student’s t-test with Welch’s correction. All data are presented as mean +/-SEM. with varying levels of significance assessed as p<0.05 (*), p<0.01 (**), p<0.001 (***), p<0.0001 (****), ns=not significant. See Table 1-1 for additional statistical details and values.

## AUTHOR CONTRIBUTIONS

The authors declare no competing interests. S.P., P.G., and D.D. conceived and designed the study. All experiments were performed by S.P. and P.G. D.M. and B.G. communicated key observations related to *smox-RNAi* and *stathmin* in the ATI background based on their own unpublished studies. The manuscript was written by S.P., P.G., and D.D. with feedback from D.M. and B.G.

## ACKNOWLEDGEMENTS

We thank Naoki Yamanaka (UC Riverside, CA, USA) and Mike O’Connor (U of Minnesota, MN, USA) who sent us other RNAi lines to test for the ATI model. We also thank the Bloomington *Drosophila* Stock Center and Developmental Studies Hybridoma Bank for additional stocks and reagents (National Institutes of Health grant P40OD018537). This study was supported by a grant from the National Institute of Neurological Disease and Stroke to D.D. (NS111414).

## Supplementary information

**Figure 4-1:**
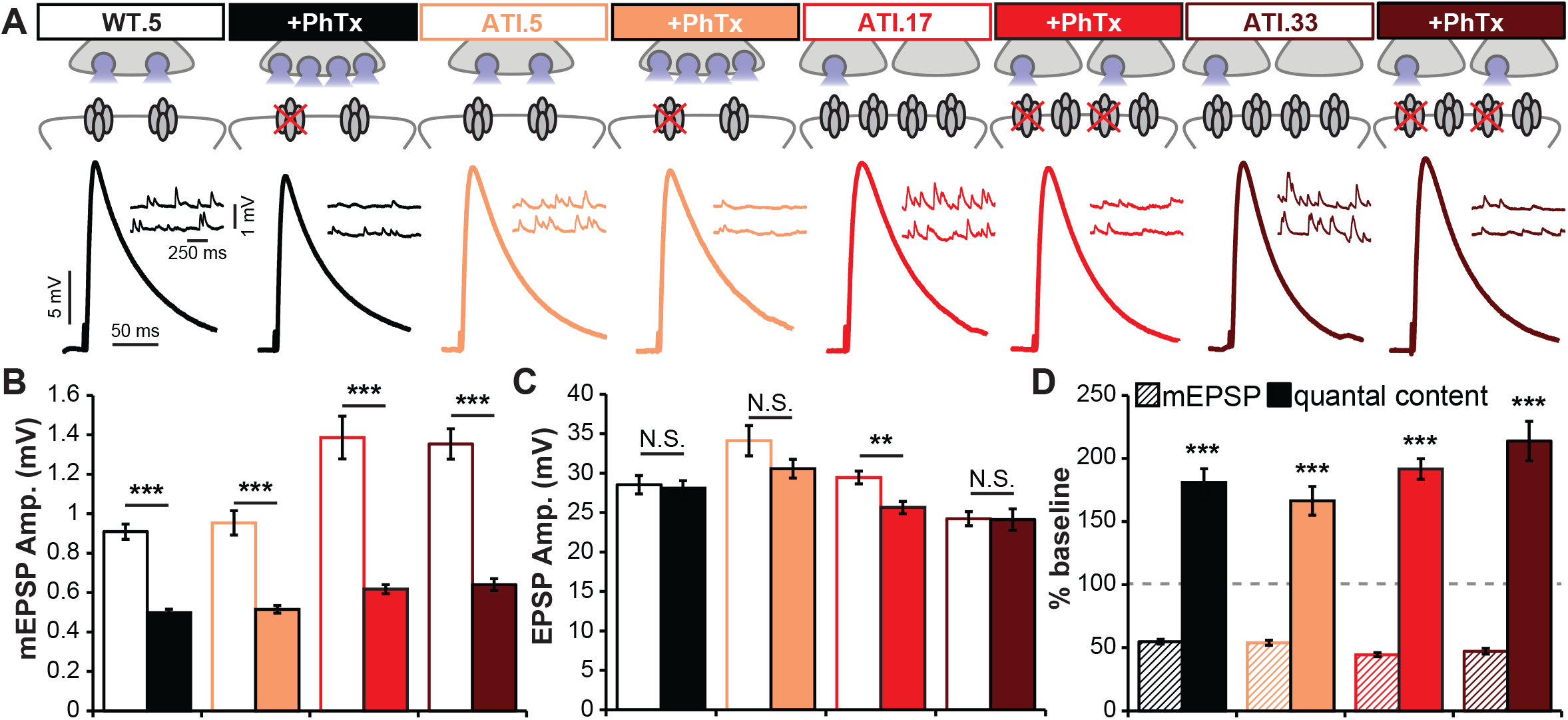
PHP can be induced and expressed across the ATI lifespan. **(A)** (Top) Schematic illustrating baseline and +PhTx conditions at NMJs for each genotype. (Bottom) Representative EPSP and mEPSP traces for each genotype at baseline and +PhTx. Quantification of mEPSP amplitude **(B)**, EPSP amplitude **(C)**, and mEPSP and quantal content values following PhTx application normalized to baseline values (-PhTx) **(D)** in the indicated genotypes. Error bars indicate ±SEM. **p≤0.01; ***p≤0.001; N.S.=not significant, p>0.05. Detailed statistical information (mean values, SEM, n, p) is shown in Table 1-1.

**Table 1-1.** Data and statistical information: All absolute values (mean, SEM, N) and statistical information (tests used and P-values) for all data points shown in Figures 1-5 and Figure 4-1 as well as supplemental data relating to these figures is shown.

**Figure 1:**
| Symbol | Genotype | Time Point days AEL | Muscle surface area $\mu\text{m}^2$ (± SEM, N) | Input resistance MΩ (± SEM, N) | EPSP amplitude mV (± SEM, N) |
| --- | --- | --- | --- | --- | --- |
| WT.5 | *W<sup>1118</sup>* | 5 | 54187.28<br>(2327.86, 10) | 6.11 (0.31, 13) | 30.25<br>(0.87, 15) |
| ATI.5 | *UAS-smox-RNAi/+; phm-Gal4/+* | 5 | 49086.17<br>(2581.39, 10) | 6 (0.32, 10) | 29.71<br>(0.95, 11) |
| ATI.9 | *UAS-smox-RNAi/+; phm-Gal4/+* | 9 | 63157.11<br>(3264.28, 10) |  |  |
| ATI.13 | *UAS-smox-RNAi/+; phm-Gal4/+* | 13 | 64947.26<br>(1929.30, 9) | 4.33 (0.42, 6) | 29.36<br>(1.59, 6) |
| ATI.17 | *UAS-smox-RNAi/+; phm-Gal4/+* | 17 | 75962.95<br>(2756.79, 10) | 2.41 (0.35, 19) | 31.03<br>(0.99, 21) |
| ATI.21 | *UAS-smox-RNAi/+; phm-Gal4/+* | 21 | 73533.58<br>(5041.38, 10) | 2.16 (0.19, 9) | 27.08<br>(1.91, 9) |
| ATI.25 | *UAS-smox-RNAi/+; phm-Gal4/+* | 25 | 65459.34<br>(4755.70, 10) |  |  |
| ATI.29 | *UAS-smox-RNAi/+; phm-Gal4/+* | 29 | 60284.69<br>(1918.83, 10) |  |  |
| ATI.33 | *UAS-smox-RNAi/+; phm-Gal4/+* | 33 | 62156.54<br>(3388.91, 10) | 3.25 (0.21, 16) | 28.89<br>(0.89, 19) |

Statistical analysis: 1-way ANOVA
| Arrays | P-value | Symbol |
| --- | --- | --- |
| WT.5 EPSP | 0.309981 | N.S. |
| ATI.5 EPSP |  |  |
| ATI.17 EPSP |  |  |
| ATI.33 EPSP |  |  |

**Figure 2:**
| Symbol | Neuronal S.A. (M4-Ib) $\mu\text{m}^2$ (± SEM) | Boutons M4-Ib # (± SEM) | BRP puncta # (± SEM) | BRP puncta size $\mu\text{m}^2$ (± SEM) | BRP puncta mean intensity % WT.5 (± SEM) | N |
| --- | --- | --- | --- | --- | --- | --- |
| WT.5 | 238.18 (10.03) | 47.54 (2.44) | 370.11<br>(12.01) | 0.089 (0.0037) | 100 (2.47) | 28 |
| ATI.5 | 237.07 (10.65) | 54.7 (3.9) | 377.8 (13.63) | 0.084 (0.0022) | 105.07 (1.89) | 10 |
| ATI.9 | 380.93 (24.62) | 78.6 (6.97) | 628.8 (28.77) | 0.085 (0.0022) | 93.12 (2.61) | 10 |
| ATI.13 | 367.88 (25.54) | 88.2 (5.89) | 681.71<br>(29.02) | 0.07 (0.0031) | 91.85 (1.72) | 10 |
| ATI.17 | 446.01 (22.63) | 106.2 (10.72) | 857.5 (37.7) | 0.12 (0.0067) | 134.44 (6.19) | 10 |
| ATI.21 | 545.13 (38.91) | 108.4 (6.06) | 991.7 (65.04) | 0.103 (0.0102) | 117.7 (9.98) | 10 |
| ATI.25 | 566.35 (53.63) | 101.2 (4.77) | 1042.8<br>(30.09) | 0.108 (0.0047) | 115.21 (3.98) | 10 |
| ATI.29 | 529.16 (35.02) | 104.6 (10.61) | 903.3 (40.15) | 0.059 (0.006) | 84.37 (3.07) | 10 |
| ATI.33 | 552.775 (20.11) | 105.4 (7.75) | 843.1 (39.06) | 0.038 (0.0074) | 73.69 (4.27) | 10 |

**Figure 3:**
| Symbol | GluRIID puncta M4-Ib<br># (± SEM) | Puncta sum intensity<br>(IIA, IIB, IID)<br>% WT.5 (± SEM) | N | mEPSP Amp.<br>mV (± SEM, N) | mEPSP freq.<br>Hz (± SEM, N) |
| --- | --- | --- | --- | --- | --- |
| WT.5 | 299.43 (11.13) | 100 (2.84)<br>100 (1.97)<br>100 (4.01) | 40 | 0.9308 (0.0411, 11) | 2.857 (0.1437, 14) |
| ATI.5 | 325.95 (23.8) | 80.13 (8.72)<br>83.49 (3.49)<br>98.75 (14.29) | 20 | 0.8799 (0.0276, 10) | 3.056 (0.1818, 9) |
| ATI.17 | 761.05 (30.61) | 149.8 (10.97)<br>143.46 (9.38)<br>132.59 (10.85) | 20 | 1.3285 (0.0657, 12) | 3.851 (0.1824, 12) |
| ATI.33 | 801.64 (25.18) | 113.09 (7.1)<br>130.63 (9.94)<br>118.81 (10.26) | 22 | 1.47 (0.0664, 10) | 3.803 (0.1845, 15) |

**Statistical Analysis: 2-tailed Student's T-test**
| Array 1 | Array 2 | P-value | Symbol |
| --- | --- | --- | --- |
| WT.5 GluRIID # | ATI.5 GluRIID # | 0.253233954 | N.S. |
| WT.5 GluRIID # | ATI.17 GluRIID # | 1.36825E-24 | *** |
| WT.5 GluRIID # | ATI.33 GluRIID # | 2.3473E-29 | *** |
| WT.5 GluRIIA intensity | ATI.5 GluRIIA intensity | 0.008809949 | ** |
| WT.5 GluRIIA intensity | ATI.17 GluRIIA intensity | 3.71503E-07 | *** |
| WT.5 GluRIIA intensity | ATI.33 GluRIIA intensity | 0.048327587 | * |
| WT.5 GluRIIB intensity | ATI.5 GluRIIB intensity | 4.08533E-05 | *** |
| WT.5 GluRIIB intensity | ATI.17 GluRIIB intensity | 1.00443E-07 | *** |
| WT.5 GluRIIB intensity | ATI.33 GluRIIB intensity | 0.000222584 | *** |
| WT.5 GluRIID intensity | ATI.5 GluRIID intensity | 0.91384072 | N.S. |
| WT.5 GluRIID intensity | ATI.17 GluRIID intensity | 0.001117841 | ** |
| WT.5 GluRIID intensity | ATI.33 GluRIID intensity | 0.047545985 | * |
| WT.5 mEPSP amplitude | ATI.5 mEPSP amplitude | 0.327204201 | N.S. |
| WT.5 mEPSP amplitude | ATI.17 mEPSP amplitude | 4.01613E-05 | *** |
| WT.5 mEPSP amplitude | ATI.33 mEPSP amplitude | 1.05448E-06 | *** |
| WT.5 mEPSP frequency | ATI.5 mEPSP frequency | 0.397753778 | N.S. |
| WT.5 mEPSP frequency | ATI.17 mEPSP frequency | 0.000225276 | *** |
| WT.5 mEPSP frequency | ATI.33 mEPSP frequency | 0.000437242 | *** |

**Figure 4:**
| Symbol | Quantal Content<br>(± SEM, N) | Failures<br>% (± SEM, N) | PPF<br>EPSC1/2 % (± SEM, N) | PPD<br>EPSC1/2 % (± SEM, N) |
| --- | --- | --- | --- | --- |
| **WT.5** | 32.02 (1.7, 11) | 51.25 (6.09, 16) | 149.91 (9.65, 12) | 74.35 (5.55, 11) |
| **ATI.5** | 34.54 (1.62, 10) | 61.39 (3.41, 9) | 158.91 (13.97, 9) | 81.34 (7.92, 8) |
| **ATI.17** | 22.38 (1.55, 12) | 79.17 (2.54, 12) | 191.04 (11.11, 14) | 96.76 (6.22, 14) |
| **ATI.33** | 18.85 (1.43, 10) | 76.73 (3.47, 13) | 178.55 (10.86, 14) | 102.09 (7.01, 13) |

**Statistical Analysis: 2-tailed Student's T-test**
| Array 1 | Array 2 | P-value | Symbol |
| --- | --- | --- | --- |
| WT.5 quantal content | ATI.5 quantal content | 0.298373008 | N.S. |
| WT.5 quantal content | ATI.17 quantal content | 0.00026744 | *** |
| WT.5 quantal content | ATI.33 quantal content | 1.18278E-05 | **** |
| WT.5 failures | ATI.5 failures | 0.249077828 | N.S. |
| WT.5 failures | ATI.17 failures | 0.000840065 | *** |
| WT.5 failures | ATI.33 failures | 0.002029811 | ** |
| WT.5 PPF | ATI.5 PPF | 0.213347891 | N.S. |
| WT.5 PPF | ATI.17 PPF | 0.003167219 | ** |
| WT.5 PPF | ATI.33 PPF | 0.006518763 | ** |
| WT.5 PPD | ATI.5 PPD | 0.198726734 | N.S. |
| WT.5 PPD | ATI.17 PPD | 0.003167234 | ** |
| WT.5 PPD | ATI.33 PPD | 0.002695262 | ** |

**Figure 5.**
| Symbol | Genotype | Time Point<br>days<br>AEL | Retractions ($\geq 1$<br>footprint)<br>% NMJs (± SEM, N) | Boutons A4-M4-Ib<br># (± SEM, N) | BRP puncta<br># (± SEM, N) |
| --- | --- | --- | --- | --- | --- |
| WT.5 | *W<sup>1118</sup>* | 5 | 8.33 (2.95, 9) |  |  |
| ATI.5 | *UAS-smox-RNAi/+; phm-Gal4/+* | 5 | 4.69 (2.29, 8) | 53.8 (7.05, 5) | 358.53<br>(10.06, 15) |
| ATI.13 | *UAS-smox-RNAi/+; phm-Gal4/+* | 13 | 5.13 (2.51, 8) | 74.71 (5.37, 7) | 631.71<br>(26.27, 7) |
| ATI.17 | *UAS-smox-RNAi/+; phm-Gal4/+* | 17 | 6.55 (4.36, 6) |  |  |
| ATI.21 | *UAS-smox-RNAi/+; phm-Gal4/+* | 21 | 6.94 (3.03, 9) |  |  |
| ATI.33 | *UAS-smox-RNAi/+; phm-Gal4/+* | 33 | 10.42 (3.84, 6) |  |  |
| stai.5 | *staiB200* | 5 | 36.31 (8.58, 11) | 9.13 (3.85, 8) | 32.63<br>(11.3, 8) |
| stai.13 | *staiB200, UAS-smox-RNAi/staiB200;*<br>*phm-Gal4/+* | 13 | 69.29 (5.02, 10) | 16.63 (1.25, 8) | 60.38<br>(13.26, 8) |

| Symbol | Genotype | Time Point<br>days<br>AEL | Footprints/<br>NMJ (A2)<br># (± SEM, N) | Footprints/<br>NMJ (A3)<br># (± SEM, N) | Footprints/<br>NMJ (A4)<br># (± SEM, N) | Footprints/<br>NMJ (A5)<br># (± SEM, N) |
| --- | --- | --- | --- | --- | --- | --- |
| stai.5 | *staiB200* | 5 | 0.14<br>(0.078, 21) | 0.29<br>(0.14, 21) | 1.09<br>(0.33, 22) | 2.10<br>(0.52, 21) |
| stai.13 | *staiB200, UAS-smox-*<br>*RNAi/staiB200; phm-Gal4/+* | 13 | 0.3<br>(0.18, 20) | 1.5<br>(0.29, 20) | 3.05<br>(0.59, 19) | 5.65<br>(0.92, 20) |

**Statistical Analysis: 2-tailed Student's T-test**
| Array 1 | Array 2 | P-value | Symbol |
| --- | --- | --- | --- |
| stai.5 Retractions | stai.13 Retractions | 0.001739115 | ** |
| ATI.5 A4 boutons | ATI.13 A4 boutons | 0.036948124 | * |
| stai.5 A4 boutons | stai.13 A4 boutons | 0.08497713 | N.S. |
| ATI.5 A4 BRP # | ATI.13 A4 BRP # | 1.55159E-10 | *** |
| stai.5 A4 BRP # | stai.13 A4 BRP # | 0.133588744 | N.S. |
| stai.5 A2 footprints | stai.13 A2 footprints | 0.418757225 | N.S. |
| stai.5 A3 footprints | stai.13 A3 footprints | 0.000400378 | *** |
| stai.5 A4 footprints | stai.13 A4 footprints | 0.004994405 | ** |
| stai.5 A5 footprints | stai.13 A5 footprints | 0.001560913 | ** |

**Figure 4-1.**
| Symbol | Baseline |  |  |  | +PhTx |  |  |  |
| --- | --- | --- | --- | --- | --- | --- | --- | --- |
| | mEPSP Amp.<br>mV (± SEM) | EPSP Amp.<br>mV (± SEM) | Quantal Content<br>(± SEM) | N | mEPSP Amp.<br>mV (± SEM) | EPSP Amp.<br>mV (± SEM) | Quantal Content<br>(± SEM) | N |
| **WT.5** | 0.909<br>(0.038) | 28.532<br>(1.166) | 31.524 (1.275) | 5 | 0.498 (0.017) | 28.097<br>(0.93) | 57.051 (3.442) | 7 |
| **ATI.5** | 0.954<br>(0.062) | 34.113<br>(1.928) | 36.154 (2.263) | 7 | 0.515 (0.019) | 30.552<br>(1.184) | 60.166 (4.105) | 7 |
| **ATI.17** | 1.386<br>(0.108) | 29.437<br>(0.81) | 22.130 (2.005) | 8 | 0.617 (0.022) | 25.643<br>(0.763) | 42.418 (1.791) | 11 |
| **ATI.33** | 1.354<br>(0.077) | 24.217<br>(0.905) | 18.063 (0.688) | 7 | 0.64 (0.03) | 24.113<br>(1.363) | 38.613 (2.820) | 10 |

**Statistical Analysis: 2-tailed Student's T-test**
| Array 1 | Array 2 | P-value | Symbol |
| --- | --- | --- | --- |
| WT.5 Baseline mEPSP | WT.5 +PhTx mEPSP | 7.74747E-07 | *** |
| ATI.5 Baseline mEPSP | ATI.5 +PhTx mEPSP | 4.8591E-06 | *** |
| ATI.17 Baseline mEPSP | ATI.17+PhTx mEPSP | 4.21332E-07 | *** |
| ATI.33 Baseline mEPSP | ATI.33 +PhTx mEPSP | 7.35985E-08 | *** |
| WT.5 Baseline EPSP | WT.5 +PhTx EPSP | 0.774221103 | N.S. |
| ATI.5 Baseline EPSP | ATI.5 +PhTx EPSP | 0.133762953 | N.S. |
| ATI.17 Baseline EPSP | ATI.17+PhTx EPSP | 0.003103703 | ** |
| ATI.33 Baseline EPSP | ATI.33 +PhTx EPSP | 0.956749384 | N.S. |
| WT.5 Baseline Quantal Content | WT.5 +PhTx Quantal Content | 0.000134197 | *** |
| ATI.5 Baseline Quantal Content | ATI.5 +PhTx Quantal Content | 0.000217442 | *** |
| ATI.17 Baseline Quantal Content | ATI.17+PhTx Quantal Content | 6.80559E-06 | *** |
| ATI.33 Baseline Quantal Content | ATI.33 +PhTx Quantal Content | 4.42488E-05 | *** |

**Table 2-1:** KEY RESOURCES TABLE.

| REAGENT/RESOURCE | SOURCE | IDENTIFIER |  |
| --- | --- | --- | --- |
| Antibodies |  |  | Dilution |
| Tetramethylrhodamine (TRITC)-conjugated phalloidin (R415) | Thermo Fisher Scientific | 41-6559-05 | 1:1000 |
| Mouse anti-Bruchpilot (nc82) | DSHB | AB_2314866 | 1:100 |
| Guinea pig anti-vGlut | (Chen et al., 2017) | N/A | 1:200 |
| Mouse anti-GluRIIA (8B4D2) | DSHB | AB_528269 | 1:50 |
| Affinity-Purified Rabbit anti-GluRIIB | (Perry et al., 2017) | N/A | 1:2000 |
| Guinea pig anti-GluRIID | (Kikuma et al., 2017) | N/A | 1:1000 |
| Rabbit anti-DLG | (Pielage et al., 2006) | N/A | 1:10000 |
| Alexa Dylite 405-conjugated secondary antibodies | Jackson ImmunoResearch Labs | 706-475-148 | 1:400 |
| Alexa Fluor 488-conjugated secondary antibodies | Jackson ImmunoResearch Labs | 706-545-148,<br>715-545-150,<br>711-545-152 | 1:400 |
| Alexa Fluor Cy3-conjugated secondary antibodies | Jackson ImmunoResearch Labs | 706-165-148,<br>715-165-150,<br>711-165-152 | 1:400 |
| Alexa Fluor 647 conjugated Goat anti-Horseradish Peroxidase | Jackson ImmunoResearch Labs | 123-605-021 | 1:200 |
| Experimental Models: Fly Lines |  |  |  |
| *UAS-smox-RNAi* | (Gibbens et al., 2011) | BDSC #41670 |  |
| *phm-Gal4* | (Ono et al., 2006) | BDSC #80577 |  |
| *staiB200* | (Graf et al., 2011) | BDSC #16165 |  |

